# A Rab11-GEF Parcas recruits Rab11 onto recycling endosomes for rhodopsin transport in *Drosophila* photoreceptors

**DOI:** 10.1101/561563

**Authors:** Yuna Otsuka, Takunori Satoh, Nozomi Nakayama, Ryota Inaba, Akiko K. Satoh

## Abstract

Rab11 and its effectors dRip11 and MyoV are essential for polarized post-Golgi vesicle trafficking to photosensitive membrane rhabdomeres in *Drosophila* photoreceptors. Here, we found that Parcas (Pcs), recently shown to have guanine-nucleotide-exchange (GEF) activity toward Rab11, co-localizes with Rab11 on the trans-side of Golgi units and post-Golgi vesicles at the base of the rhabdomeres in pupal photoreceptors. Pcs fused with the EM-tag APEX2 localizes on 150-300 nm vesicles at the trans-side of Golgi units, which are presumably fly recycling endosomes (RE). Loss of Pcs impairs Rab11 localization on the trans-side of Golgi units and induces the cytoplasmic accumulation of post-Golgi vesicles bearing rhabdomere proteins, as observed in Rab11-deficiency. In contrast, loss of the specific subunits of TRAPPII, another known Rab11-GEF, does not cause any defects on the eye development nor the transport of rhabdomere proteins, however, simultaneous loss of TRAPPII and Pcs shows severe defects on eye development. These results indicated that in pupal photoreceptors, Pcs is the predominant Rab11-GEF, and TRAPPII performs a function that is redundant but subsidiary to that of Pcs.

**Summary statement:** Parcas, a Rab11-GEF, recruits Rab11 onto Golgi-associated RE and post-Golgi vesicles in fly photoreceptors. Loss of Parcas results in rhodopsin accumulation within the cytoplasm, similar to that observed under Rab11-deficiency.

## Introduction

Sensory neurons exploit a common mechanism of polarized epithelial cell differentiation, amplification, and specialization in apical plasma membranes to build sensory organelles. An example is *Drosophila* photoreceptor morphogenesis, where a late-pupal surge of secretory membrane traffic expands the apical photosensory membrane, the rhabdomere.

Rab proteins are small GTPases that control membrane traffic and maintain distinct organelle identities. More than 60 mammalian and 31 *Drosophila* Rab proteins regulate specific transport steps and pathways (Stenmark, 2009; Welz et al., 2014; Zhang et al., 2007). In *Drosophila* photoreceptors, 3 Rab proteins sequentially regulate the transport of Rh1, the rhodopsin expressed in R1–6 outer retinal photoreceptor cells. Rab1 and Rab6 regulate Rh1 transport from the ER to the *cis-*Golgi and trans-Golgi network (TGN) to the RE, respectively (Iwanami et al., 2016; Satoh et al., 1997; Satoh et al., 2005). Rab11 and its effectors, dRip11 and MyoV, allow the Rh1-bearing post-Golgi vesicles to invade the exclusive retinal terminal web, the bundle of actin filaments which plus-ends are anchored to the microvilli base (Li et al., 2007; Satoh et al., 2005).

The activities of Rab proteins are regulated by guanine nucleotide exchange factors (GEFs) and GTPase-activating proteins (GAPs) (Barr and Lambright, 2010; Ishida et al., 2016). GEFs activate Rab proteins by exchanging a GDP for a GTP on a given Rab protein. GAPs inactivate Rab proteins by facilitating the GTPase activity of Rab proteins to complete the Rab cycle. Two GEF proteins are reported to be involved in Rh1 transport, specifically, Rich (a Rab6-GEF) and Crag (a Rab11-GEF). *Rich*-null mutant (*Rich*^*1*^) photoreceptors exhibit phenotypes similar to those exhibited by Rab6-null mutant photoreceptors (Iwanami et al., 2016); however, the phenotypes of *Crag*-null mutant (*Crag*^*CJ101*^) photoreceptors are quite different from those of Rab11-deficient photoreceptors, consistent with the idea that Crag facilitates light-dependent Rh1 transport in adult flies, but not during pupal development (Xiong et al., 2012).

Transport protein particle (TRAPP) complexes were originally identified in yeast as RabGEFs, and these proteins share a core set of subunits (Jones et al., 2000; Pinar et al., 2015; Sacher et al., 2008; Thomas and Fromme, 2016; Thomas et al., 2018). A recent study indicated that *Drosophila* possesses two TRAPP complexes (II and III), and both activate Rab1. One of these complexes, TRAPPII, also activates Rab11 (Riedel et al., 2018). Despite this, the specific subunits of TRAPPII are not essential, and one of these subunits has been shown to localize on the cis-side of Golgi rather than the trans-side of Golgi or recycling endosomes, where Rab11 is thought to be activated (Riedel et al., 2018).

REI-1 is another Rab11-GEF that was recently identified in *C. elegans*, and this protein does not contain any known Rab-GEF domain, yet exhibits a strong Rab11-GEF activity *in vitro* (Sakaguchi et al., 2015). *C. elegans* models of double-mutant REI-1 and its close paralog REI-2 exhibit a loss of Rab11 localization to the late-Golgi compartments, RE, and cleavage furrows, as well as delayed cytokinesis (Sato et al., 2008). Both *Drosophila* Rab11 and the REI-1 ortholog parcas/poirot (Pcs/Prt) are necessary for the specific localization of osker mRNA and Osker protein to the oocyte posterior pole, although the mechanism underlying this localization remains unknown (Jankovics et al., 2001; Sinka et al., 2002). In the Pcs-deficient fly line, a hobo insertion into the first intron of the *pcs* gene, *pcs*^*gs*^, is not lethal, and this mutant exhibits a grandchildless phenotype possibly caused by the lack of Rab11 activation in the oocyte. Pcs is also involved in muscle morphogenesis and Wnt signaling (Beckett and Baylies, 2006). Additionally, a recent study indicated that in addition to TRAPPII complex, pcs possesses GEF activity toward Rab11 *in vitro*, and *pcs*^*gs*^ causes synthetic lethality with the null mutation of TRAPPC9, a specific subunit of TRAPPII (Riedel et al., 2018). Here we investigated the function of Pcs and TRAPPII in *Drosophila* photoreceptors.

## Results and Discussion

### Pcs-deficient photoreceptors accumulate rhabdomeric proteins in the cytoplasm

Rab11 and its effectors, dRip11 and MyoV, are essential for the post-Golgi trafficking of Rh1 to the rhabdomeres, and their loss causes Rh1 accumulation within the photoreceptor cytoplasm (Li et al., 2007; Satoh et al., 2005). If Pcs functions as a Rab11-GEF in the developing photoreceptors, lack of Pcs should cause defects in Rh1 transport similar to those caused by the loss of Rab11. Thus, we investigated Rh1 localization in two hypomorphic alleles with transposon insertions, *pcs*^*GS7166*^ in the first intron, or *pcs*^*NP2623*^ in the 5’UTR. Additionally, we used the CRISPR/Cas9 system to generate a deletion mutant allele, *pcs*^*Δ1*^, that is thought to act as a *null* allele (Figure 1A). Although these three alleles are viable, we observed mosaic retinas with both the wild type and mutant photoreceptors, which allow us to assess the phenotype relative to the wild type. We found cytoplasmic accumulation of Rh1 in the late pupal retinas of *pcs*^*GS7166*^, *pcs*^*NP2623*^, and *pcs*^*Δ1*^ homozygous photoreceptors (Figures 1B, D and S1A), indicating that Pcs functions as a possible Rab11-GEF in the developing photoreceptors. The cytoplasmic accumulation of Rh1 disappeared completely in *pcs*^*GS7166 ex1*^ homozygous photoreceptors, where the P-element insertion is precisely excised from *pcs*^*GS7166*^ by the delta2-3 transposase (Figure 1C). Thus, P-element insertion into the first intron of the *pcs* gene is the sole cause of Rh1 accumulation in the *pcs*^*GS7166*^ photoreceptor cytoplasm. The quantification of the ratio of Rh1 staining in the cytoplasm against that in the whole photoreceptor that includes both rhabdomeres and cytoplasm, in the wild type, *pcs*^*GS7166*^, *pcs*^*GS7166 ex1*^, and *pcs*^*Δ1*^ alleles (Figure 1H) verified more than 80% of Rh1 staining in the cytoplasm of *pcs*^*GS7166*^ and *pcs*^*Δ1*^ homozygous photoreceptors. In *pcs*^*gs*^, a piggyBac insertion within the first intron that has been reported as a null mutation (Sinka et al., 2002), Rh1 localized to the rhabdomeres (Figure S1B), suggesting that this piggyBac insertion does not effectively reduce the expression of Pcs protein in the eyes.

**Fig 1.**
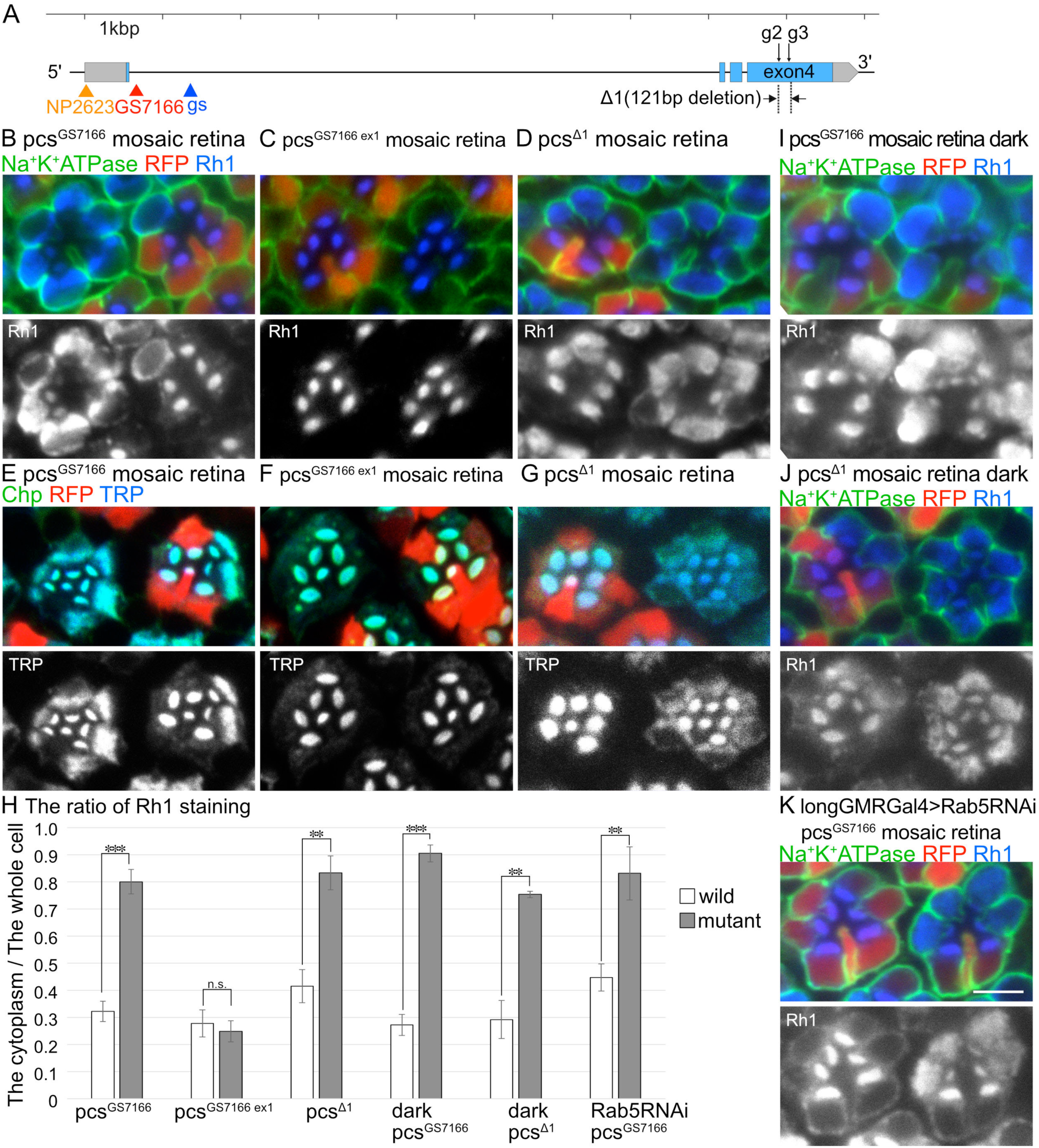
Rhabdomere proteins accumulate in the cytoplasm of Pcs-deficient photoreceptors. (A) Schematic view of *pcs* gene within the genome. Arrowheads indicated insertion sites for three transposons. g2 and g3 indicate CRISPR target sites, and *pcs*^*Δ1*^, a null allele made by the CRISPR/Cas9 system, has a 121-bp deletion between g2 and g3. Gray bars represent the mRNA sequences, and blue bars represent the coding sequences. (B-D) Immunostaining of *pcs*^*GS7166*^ (B), *pcs*^*GS7166 ex1*^ (C), or *pcs*^*Δ1*^ (D) mosaic eyes with anti-Na^+^K^+^ATPase (green) and anti-Rh1 (blue) antibodies. RFP (red) indicates wild-type cells. (E-G) Immunostaining of *pcs*^*GS7166*^ (E), *pcs*^*GS7166 ex1*^ (F), or *pcs*^*Δ1*^ (G) mosaic eyes with anti-Chp (green) and anti-TRP (blue) antibodies. RFP (red) indicates wild-type cells. (H) The ratio of signal strength for Rh1 staining in the cytoplasm against that of the whole cells was plotted. White bars indicate the wild type cells and the grey bars indicate mutant cells in *pcs*^*GS7166*^, *pcs*^*GS7166 ex1*^, *pcs*^*Δ1*^, dark-reared *pcs*^*GS7166*^, dark-reared *pcs*^*Δ1*^ mosaic retinas, and *pcs*^*GS7166*^ mosaic retinas expressing Rab5RNAi by longGMR-Gal4. More than 60 each of photoreceptors for both the wild type and mutant were measured. Error bars indicate s.d. Significance according to Student’s *t-*test: \*\*\**p* < 0.001 and \*\**p* < 0.01. (I, J) Immunostaining of dark-reared *pcs*^*GS7166*^ (I) or dark-reared *pcs*^*Δ1*^ mosaic eyes (J) with anti-Na^+^K^+^ATPase (green) and anti-Rh1 (blue) antibodies. RFP (red) indicates wild-type cells. (K) Immunostaining of *pcs*^*GS7166*^ mosaic eyes expressing Rab5RNAi by longGMR-Gal4 with anti-Na^+^K^+^ATPase (green) and anti-Rh1 (blue) antibodies. RFP (red) indicates wild-type cells. Scale bars: 5 µm (B-G, I-K).

Similar to Rab11-deficient photoreceptors, both TRP, a cation channel essential for photo-transduction (Montell and Rubin, 1989), and Chaoptin (Chp), an adhesion molecule bundling microvilli into the rhabdomeres (Reinke et al., 1988), accumulated in the cytoplasm of *pcs*^*GS7166*^ and *pcs*^*Δ1*^ photoreceptors (Figure 1E, G), but not in the cytoplasm of *pcs*^*GS7166 ex1*^ photoreceptors (Figure 1F). Thus, the transport of rhabdomeric proteins was inhibited by the loss of Pcs. Na^+^K^+^ATPase, however, localized normally in basolateral membranes in *pcs*^*GS7166*^, *pcs*^*Δ1*^, and *pcs*^*NP2623*^ homozygous photoreceptors (Figures 1B, D and S1A).

### Inhibition of Rh1-endocytosis does not compromise Rh1-accumulation caused by Pcs-deficiency

A recent study indicated that in *Drosophila* photoreceptors, dPIP4K regulates clathrin-mediated endocytosis (CME) from the plasma membrane, and its absence causes increased CME uptake, leading to an increased number of Rh1-loaded early endocytic vesicles (Kamalesh et al., 2017). This cytoplasmic accumulation of Rh1 in dPIP4K-deficient photoreceptors is light-dependent, likely because of light-stimulated endocytosis, and this phenotype is rescued by the inhibition of endocytosis by Rab5RNAi. To investigate whether Rh1 accumulation in Pcs-deficient cells is caused by increased endocytosis, we investigated Rh1 localization in dark-reared homozygous *pcs*^*GS7166*^ *and pcs*^*Δ1*^, and also in the *pcs*^*GS7166*^ mosaic retinas expressing Rab5RNAi. We found high levels of Rh1 accumulation in the cytoplasm of dark-reared *pcs*^*GS7166*^ *and pcs*^*Δ1*^ homozygous photoreceptors (Figure 1I, J) and also in the cytoplasm of *pcs*^*GS7166*^ photoreceptors expressing Rab5RNAi (K). Quantification of Rh1 accumulation indicated that the degree of Rh1 accumulation is not reduced under dark conditions compared to that observed under light conditions, and accumulation is also not affected by the expression of Rab5RNAi (Figure 1H). These results indicated that Pcs is likely to regulate biosynthetic pathways and not the endocytosis of Rh1.

### Cytoplasmic vesicle accumulation in Pcs-deficient photoreceptors

To more precisely compare the phenotypes between Pcs-deficiency and the loss of the Rab11/dRip11/MyoV complex, we observed thin sections of wild type *w*^*1118*^, *pcs*^*NP2623*^, *pcs*^*GS7166*^, *pcs*^*GS7166 ex1*^, and *pcs*^*Δ1*^ pupal photoreceptors using electron microscopy (Figures 2A-H and S1C, D). We found smaller rhabdomeres and cytoplasmic vesicle accumulations in *pcs*^*GS7166*^, *pcs*^*NP2623*^, and *pcs*^*Δ1*^ photoreceptors, while the wild type, *w*^*1118*^, and *pcs*^*GS7166 ex1*^ photoreceptors remained unchanged. The appearance of these vesicles resembled the vesicles that accumulated upon deficiency of the Rab11, dRip11, and MyoV complex (Li et al., 2007; Satoh et al., 2005). We measured the area of cytoplasmic vesicles in wild type, *pcs*^*GS7166*^, *pcs*^*GS7166 ex1*^, and *pcs*^*Δ1*^ pupal photoreceptors, and found that the areas occupied by the cytoplasmic vesicles were significantly increased in *pcs*^*GS7166*^ and *pcs*^*Δ1*^ pupal photoreceptors (Figure 2I). The cisternae of Golgi units appeared slightly swollen, but they still stacked well (Figure 2G, H arrows). Mitochondria and ER were normal in *pcs*^*GS7166*^ and *pcs*^*Δ1*^ photoreceptors (Figure 2G, H asterisks and arrowheads).

**Fig 2.**
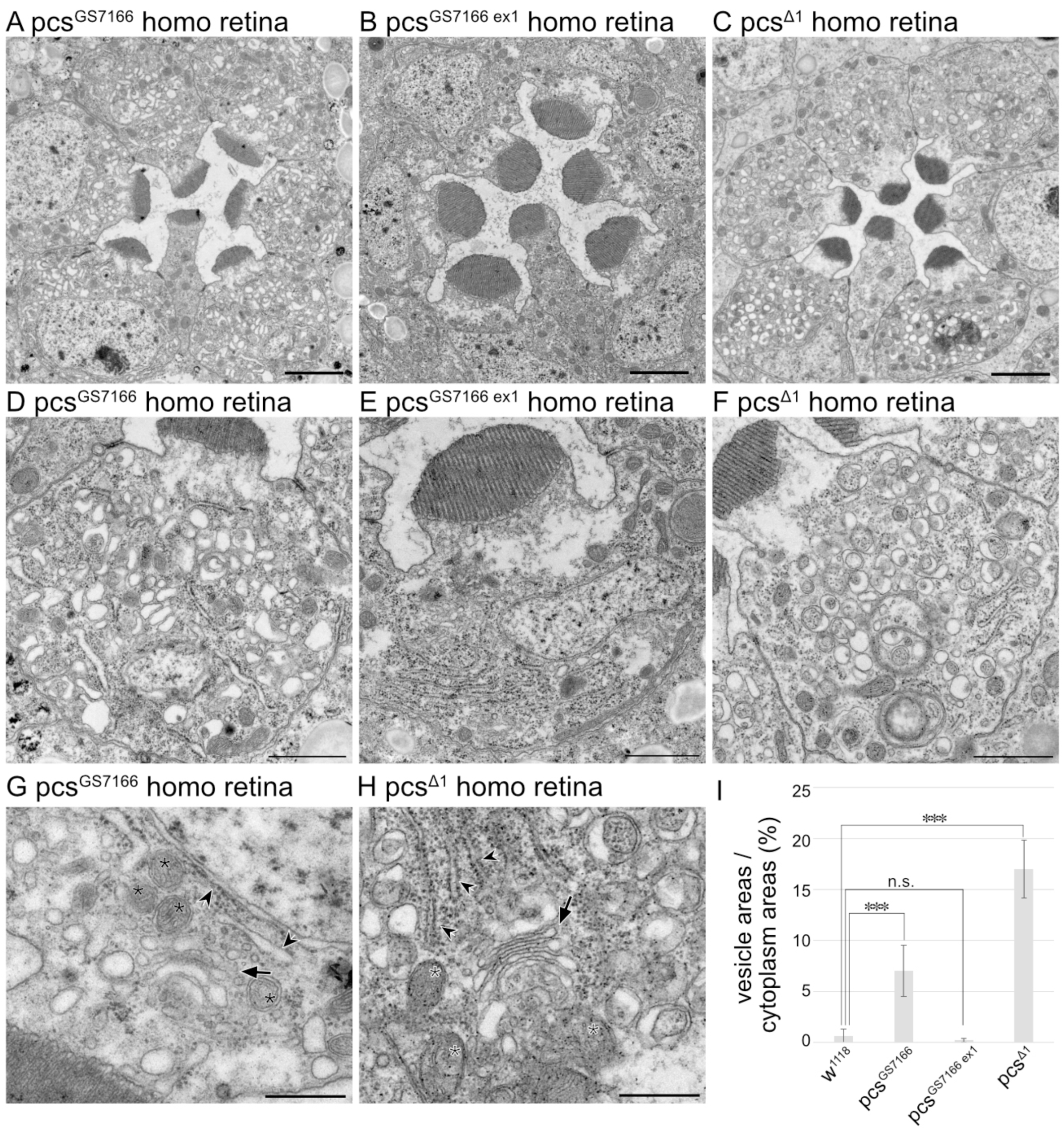
Rhabdomere proteins accumulate within the cytoplasm of Pcs-deficient photoreceptors. (A-C) Electron micrographs of the ommatidia obtained from homozygous flies of *pcs*^*GS7166*^ (A), *pcs*^*GS7166 ex1*^ (B), and *pcs*^*Δ1*^ (C). (D-F) Electron micrographs of the photoreceptor obtained from homozygous flies of *pcs*^*GS7166*^ (D), *pcs*^*GS7166 ex1*^ (E), and *pcs*^*Δ1*^ (F). (G, H) Magnified electron micrographs of *pcs*^*GS7166*^ (G) and *pcs*^*Δ1*^ (H). (I) Percentage of the area of vesicles versus the area of the cytoplasm of the wild-type (*w* ^*1118*^), *pcs*^*GS7166*^, *pcs*^*GS7166 ex1*^, and *pcs*^*Δ1*^. More than 15 photoreceptors were measured for each genotype. Error bars indicate s.d. Significance according to Student’s *t-*test: \*\*\**p* < 0.001. Scale bars: 2 µm (A–C), 1 µm (D–F), 1 µm (G, H).

### Pcs co-localizes with Rab11 on the RE and post-Golgi vesicles at the rhabdomere base

Rab11 localizes to the Golgi-associated RE and the post-Golgi vesicles at the rhabdomere base (Iwanami et al., 2016; Satoh et al., 2005). We next compared the localization of Pcs and Rab11. We failed to detect endogenous Pcs proteins using anti-Pcs antibodies, which were kindly gifted by Dr. Baylies and Dr. Erdelyi (Beckett and Baylies, 2006; Sinka et al., 2002). Therefore, we created transgenic flies expressing UAS-V5::Pcs, and expressed V5::Pcs in late pupal photoreceptors using Rh1-Gal4. V5::Pcs co-localized with Rab11 both in the cytoplasmic puncta (Figures 3A arrowheads), which are presumably RE associated with the trans-side of Golgi units, and in the puncta at the base of the rhabdomeres (Figures 3A arrows), which are the post-Golgi vesicles bearing newly synthesized rhabdomere proteins (Figures 3A, B). Line plots of fluorescent intensities of anti-V5 and anti-Rab11 (Figure 3C) through the perinuclear region from a Golgi unit to the rhabdomere (arrow in Figure 3B) indicated that the V5::Pcs signal at the RE was greater than the V5::Pcs signal at the base of the rhabdomeres, although the Rab11 signal detected at the base of the rhabdomere was greater than that observed at the RE.

**Fig 3.**
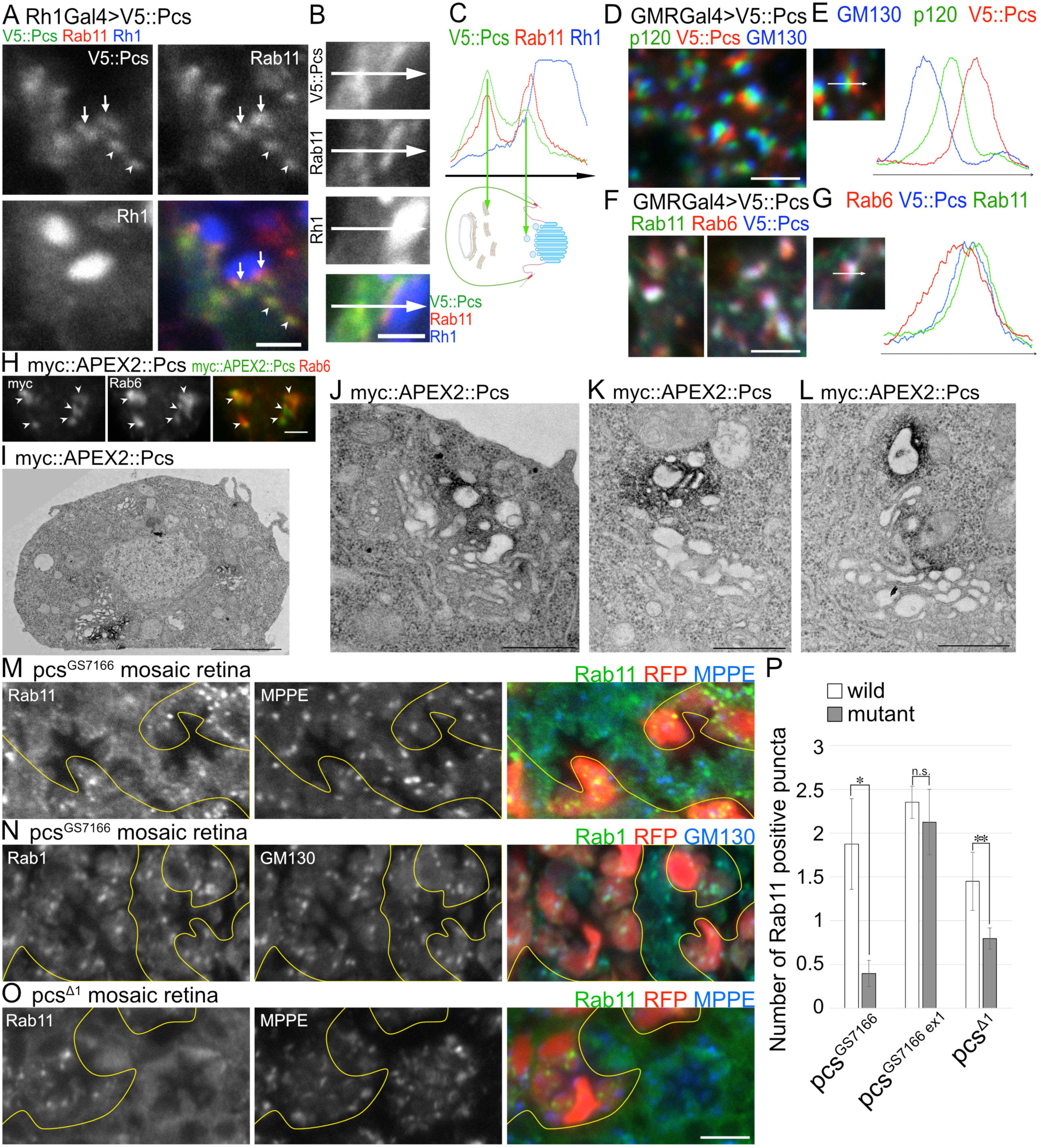
V5::Pcs co-localizes with Rab11 on RE and post-Golgi vesicles at the rhabdomere base. (A, B) Immunostaining of the wild-type photoreceptors expressing V5::Pcs driven by Rh1-Gal4 stained with anti-V5 (green), anti-Rab11 (red), and anti-Rh1 (blue) antibodies. (C) Plots of signal intensities and the distance in µm on the arrow in (B) show the overlap between channels. Respective signals from V5 (green), Rab11 (red), and Rh1 (blue) antibodies. (D) Immunostaining of wild-type eye expressing V5::Pcs driven by GMR-Gal4 stained with anti-GM130 (blue), anti-p120 (green), and anti-V5 (red) antibodies. (E) Plots of signal intensities and the distance in µm on the arrow show the overlap between channels. Respective signals from GM130 (blue), anti-p120 (green), and anti-V5 (red) antibodies. (F) Immunostaining of wild-type eye expressing V5::Pcs driven by GMR-Gal4 stained with anti-Rab11 (green), anti-Rab6 (red), and anti-V5 (blue) antibodies. (G) Plots of signal intensities and the distance in µm on the arrow show the overlap between channels. Respective signals from anti-Rab11 (green), anti-Rab6 (red), and anti-V5 (blue) antibodies. (H) Immunostaining of S2 cells expressing myc::APEX2::Pcs with anti-myc (green) and anti-Rab6 (Red) antibodies. (I-L) Electron micrographs of S2 cells expressing myc::APEX2::Pcs with anti-myc. APEX2 localization is visualized by osmium-stained DAB polymer. (M-O) Immunostaining of *pcs*^*GS7166*^ (M, N) or *pcs*^*Δ1*^ (O) mosaic eyes with anti-Rab11 (green) and anti-MPPE (blue) antibodies (M, O), or anti-Rab1 (green) and anti-GM130 (blue) antibodies (N). RFP (red) indicates wild-type cells. (P) Plots of the Rab11 puncta numbers in the cytoplasm of the wild type (white bars) and mutant cells (grey bars) in *pcs*^*GS7166*^, *pcs*^*GS7166 ex1*^, and *pcs*^*Δ1*^ mosaic retina. More than 60 each of photoreceptors for both the wild type and mutant were measured. Error bars indicate s.d. Significance according to Student’s *t-*test: \*\**p* < 0.01 and \**p* < 0.05. Scale bars: 2 µm (A, B), 5 µm (D, F), 2 µm (H), 2 µm (I), 500 nm (J-L), 5 µm (M-O).

We next investigated the detailed localization of V5::Pcs to Golgi units using young pupal retina, a tissue that possesses well-developed Golgi units. V5::Pcs signals were localized to the trans-side of the Golgi units, and separated from both the cis-Golgi marker GM130 and medial-Golgi marker p120 (Figure 3D, E). V5::Pcs, however, co-localized with Rab11 at the RE and extended toward the TGN, where Rab6 localizes (Figure 3F, G). Thus, Pcs co-localized with Rab11, but Pcs localization was shifted slightly away from Rab11 and toward the upstream TGN in the polarized transport pathway to the rhabdomere. These localization studies suggested that Pcs works upstream of Rab11, and this is in agreement with the idea that Pcs works as a Rab11-GEF.

To investigate Pcs localization using electron micrograph, we employed a recently developed genetic EM-tag, APEX2, that catalyzes the polymerization and local deposition of DAB, which in turn provides EM contrast after treatment with OsO_4_ (Lam et al., 2015; Martell et al., 2012). First we confirmed that myc::APEX2::Pcs associates with Golgi units in S2 cells by immunostaining using anti-myc (Figure 3H). In the electron micrographs of S2 cells expressing myc::APEX2::Pcs, electron dense DAB staining was found on the trans-side of Golgi units (Figure 3I, J) or at some distance from the trans-side of Golgi units (Figure 3K, L). DAB staining of myc::APEX2::Pcs primarily associated with 150-300 nm distorted-vesicles or cisternae (Figure 3J-L). These observations suggest that these membrane structures are likely to represent recycling endosomes in *Drosophila*.

### Pcs is necessary for Rab11 recruitment to the RE and post-Golgi vesicles

We next investigated whether Pcs is required for Rab11 localization to the Golgi-associated RE and post-Golgi vesicles by immunostaining of *pcs*-deficient mosaic retina (Figures 3M, O and S1E). The strong punctate staining of Rab11 seen in the wild type photoreceptors was absent in *pcs*^*GS7166*^ and *pcs*^*Δ1*^ homozygous photoreceptors. As there was no significant difference in the staining of the medial-Golgi marker MPPE between the wild type and homozygous *pcs*^*GS7166*^ and *pcs*^*Δ1*^ photoreceptors, the Golgi units themselves were not eliminated in *pcs*-deficient photoreceptors, consistent with our observations by electron microscopy (Figure 2G, H arrows). Rab11-localization to Golgi-associated RE and post-Golgi vesicles in *pcs*^*GS7166*^ was also rescued in *pcs*^*GS7166 ex1*^ photoreceptors, indicating that Pcs function is essential for Rab11 localization to these compartments (Figure S1F, H). We also investigated Rab1 localization in *pcs*^*GS7166*^ mosaic retina (Figure 3N). In contrast to Rab11, Rab1 localized normally to Golgi units with GM130, indicating that Pcs is involved in the recruitment of Rab11 but not of Rab1. Thus, Rab11 recruitment to Golgi-associated RE and post-Golgi vesicles is dependent upon Pcs, indicating that Pcs works as a Rab11-GEF in fly photoreceptors.

### Pcs is not necessary for Rab11 membrane recruitment

As Rab-GEFs regulate the recruitment of their target Rab proteins to the specific organelle membrane, we compared the relative amount of Rab11 on membrane fractions in *w*^*1118*^, *pcs*^*GS7166*^, *pcs*^*GS7166 ex1*^, *w*^*1118*^/deletion, and *pcs*^*NP2623*^/deletion fly heads. In every genotype test, 52-60% of Rab11 was found in the membrane fraction with no significant differences between the wild type (*w*^*1118*^, *pcs*^*GS7166 ex1*^, *w*^*1118*^/deletion) and pcs-deficient (*pcs*^*GS7166*^, *pcs*^*NP2623*^/deletion) heads (Figures S1G, I). We also confirmed that the total amount of Rab11 was unchanged in these flies (Figures S1G, I), similar to observations from a previous report (Sakaguchi et al., 2015). Thus, Pcs is necessary for the specific localization of Rab11 to the trans-side of Golgi units and post-Golgi vesicles (Figure 3M and S1G), but the protein is not essential for membrane recruitment (Figures S1H, J).

### TRAPPIII, but not TRAPPII, is necessary for Rh1 transport

A recent study indicated that both *Drosophila* TRAPPIII and TRAPPII complexes activate Rab1, and TRAPPII also activates Rab11 in a biochemical GEF-assay (Riedel et al., 2018). Given this, we investigated the functions of TRAPPIII and TRAPPII on Rh1 transport. TRAPPIII and TRAPPII share 7 subunits (TRAPPC1, C2, C2L, C3, C4, C5 and C6. TRAPPC8, C11, C12, and C13 are TRAPPIII-specific and TRAPPC9 and C10 are TRAPPII-specific subunits (Riedel et al., 2018). We first investigated the functions of the shared core subunits, TRAPPC2 (Trs20) and TRAPPC6 (Trs33), and the TRAPPIII-specific subunits, TRAPPC8 (Trs85) and TRAPPC11 (gryzun), as their null insertional mutations, *TRAPP2C*^*c00766*^, *TRAPPC6*^*f00985*^, *TRAPPC8*^*L3809*^, and *TRAPPC11*^*MB06920*^, are available publicly. As all of the insertional mutations are lethal, we combined them with corresponding FRTs and investigated the resulting mosaic retinas containing both the wild type and null homozygous photoreceptors. We found that Rh1 accumulation to the rhabdomeres was completely inhibited in *TRAPP2C*^*c00766*^, *TRAPPC6*^*f00985*^, *TRAPPC8*^*L3809*^, and *TRAPPC11*^*MB06920*^ homozygous photoreceptors (Figure 4A, B and Figure S3A, B). In contrast to that under Rab11- or Pcs-deficiency, Rh1 was not accumulated in the cytoplasm, but instead became undetectable. The immunostaining of Na^+^K^+^ATPase at the basolateral membrane was also reduced. These phenotypes were similar to those caused by the loss of Syx5, which regulates ER to Golgi transport, and these findings are in agreement with those of a recent study that showed that fly TRAPPIII is a Rab1-GEF (Riedel et al., 2018; Satoh et al., 2016). We next investigated the influence of TRAPPIII-deficiency on Rab1 and Rab11 localization. In both *TRAPPC6*^*f00985*^ and *TRAPPC8*^*L3809*^ homozygous photoreceptors, the punctate staining of Rab1 on the cis-Golgi was undetectable, while Rab11 foci were still clearly detected. Punctate staining of the Golgi markers GM130 and MPPE was also significantly reduced (Figure 4C-F). Thus, TRAPPIII is likely to function as a Rab1-GEF, and this complex is necessary for Rh1 transport, presumably from ER to cis-Golgi in fly photoreceptors.

**Fig 4.**
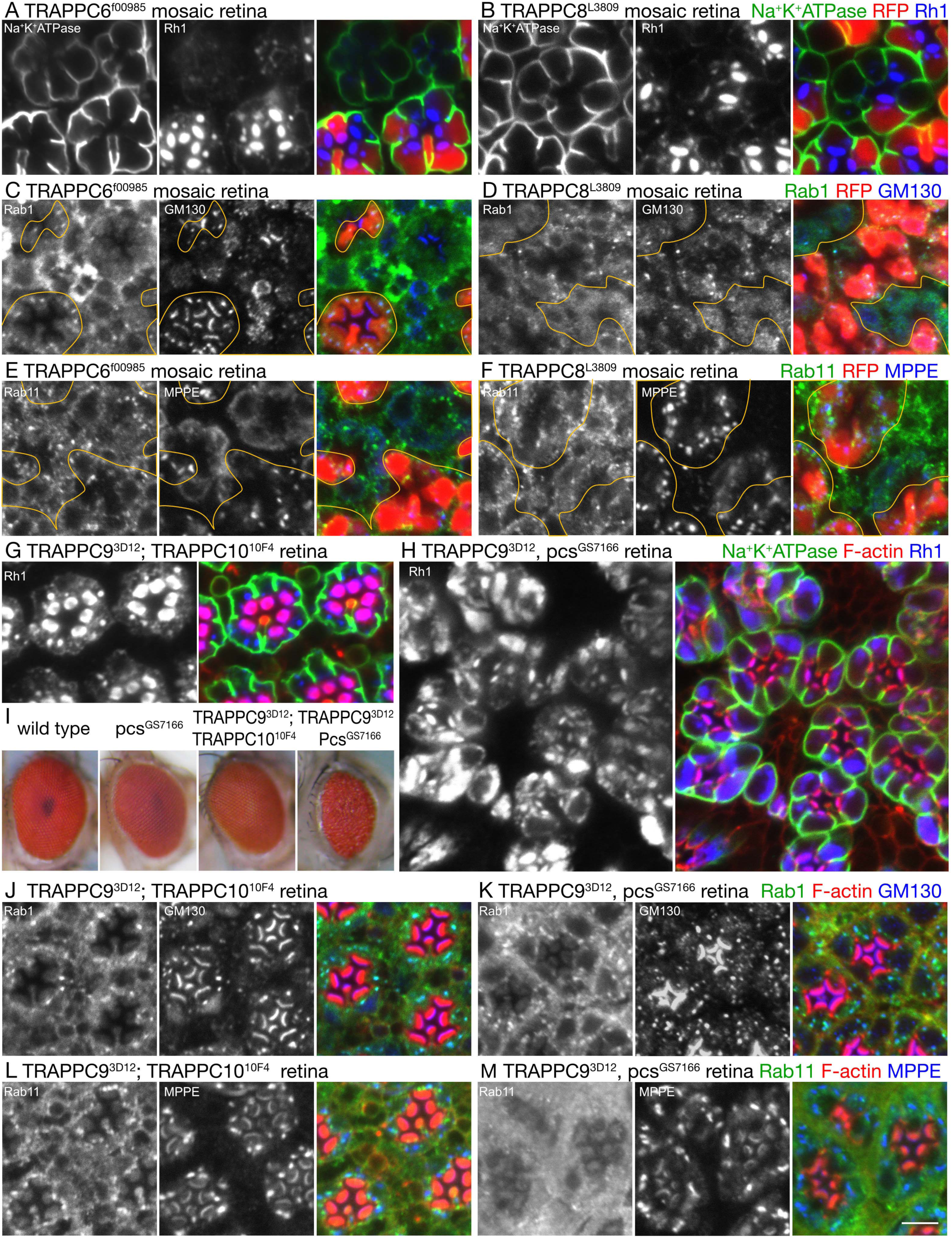
Rh1 transport is inhibited in TRAPPIII-deficient photoreceptors. (A-F) Immunostaining of *TRAPPC6*^*f00985*^ (A, C, E) or *TRAPPC8*^*L3809*^ mosaic eyes (B, D, F) with anti-Na^+^K^+^ATPase (green) and anti-Rh1 (blue) antibodies (A, B), anti-Rab1 (green) and anti-GM130 (blue) antibodies (C, D), or anti-Rab11 (green) and anti-MPPE (blue) antibodies (E, F). RFP (red) indicates wild-type cells. (G, H) Immunostaining of *TRAPPC9*^*3D12*^; *TRAPPC10*^*10F4*^ (G), or *TRAPPC9*^*3D12*^, *pcs* ^*GS7166*^ retinas (H) with anti-Na^+^K^+^ATPase (green), anti-Rh1 (blue) antibodies, and phalloidin (red). (I) Eyes of the wild type, *pcs*^*GS7166*^, *TRAPPC9*^*3D12*^; *TRAPPC10*^*10F4*^, and *TRAPPC9*^*3D12*^, *pcs*^*GS7166*^ are observed by stereomicroscopy. (J-M) Immunostaining of *TRAPPC9*^*3D12*^; *TRAPPC10*^*10F4*^ (J, L), or *TRAPPC9*^*3D12*^, *pcs*^*GS7166*^ retinas (K, M) with anti-Rab1 (green), anti-GM130 (blue) antibodies, and phalloidin (red) (J, K), or with anti-Rab11 (green), anti-MPPE (blue) antibodies and phalloidin (red) (L, M). Scale bars: 5 µm (A–H, J–M).

We made deletion mutants for the TRAPPII-specific subunits TRAPPC9 and C10 by imprecise excisions of P-elements. Both deletion mutants, *TRAPPC9*^*3D12*^ and *TRAPPC10*^*10F4*^, are thought to be null; however, both were homozygous-viable, as reported previously (Riedel et al., 2018), and no phenotype was observed on their retinas (data not shown). Even *TRAPPC9*^*3D12*^; *TRAPPC10*^*10F4*^ double-mutant flies were viable, and Rh1 and Na^+^K^+^ATPase normally localized on rhabdomeres and the basolateral membranes of their photoreceptors (Figure 4G). Additionally, Rab1 and Rab11 also localized normally to the Golgi units (Figure 4J, L). Conversely, the *TRAPP9*^*3D12*^, *pcs*^*GS7166*^ double-homozygous mutant exhibited synthetic lethality, and this is in agreement with the results of a previous study (Riedel et al., 2018). The lethal stage of *TRAPP9*^*3D12*^, *pcs*^*GS7166*^ double-homozygous mutant flies occurred immediately prior to emersion. Interestingly, the retinas of *TRAPP9*^*3D12*^, *pcs*^*GS7166*^ double-homozygous mutant pupae were small and exhibited severe roughness (Figure 4I). We dissected these small eyes and investigated the localization of Rh1 and Na^+^K^+^ATPase. As expected from the rough eye phenotype, ommatidia were not well aligned and vast spaces were found between the ommatidia in *TRAPP9*^*3D12*^, *pcs*^*GS7166*^ retinas (Figure 4H). Thus, the developmental processes of retinas were severely affected by simultaneous loss of TRAPPII and Pcs. Conversely, Rh1 transport defects of *TRAPP9*^*3D12*^, *pcs*^*GS7166*^ double-homozygous mutant photoreceptors were not significantly worse than those of *pcs*^*GS7166*^ single-homozygous mutant photoreceptors, as the majority of Rh1 was not detected on the rhabdomeres and instead accumulated in the cytoplasm (Figure 4H), similar to that in *pcs*^*GS7166*^ single-mutant photoreceptors. Na^+^K^+^ATPase localized normally at the basolateral membrane, and Rab1 localized normally on Golgi units with GM130, but Rab11 was largely diffused in TRAPP9^3D12^, pcs^GS7166^ photoreceptors (Figure 4K, M). These results indicate that in the pupal photoreceptors, Pcs functions as the predominant Rab11-GEF, while TRAPPII functions as a subsidiary Rab11-GEF that can partially compensate for Pcs-deficiency.

## Materials and Methods

### *Drosophila* stocks and genetics

Flies were grown at 20–25°C on standard cornmeal–glucose–agar–yeast food unless indicated otherwise. Carotenoid-deprived food was prepared from 1% agarose, 10% dry-yeast, 10% sucrose, 0.02% cholesterol, 0.5% propionate, and 0.05% methyl 4-hydroxybenzoate.

The fly stocks obtained from the Bloomington Drosophila Stock Center (BDSC), the Kyoto Drosophila Genomics and Genetic Resource (DGGR) and Exelixis at Harvard medical School were referred to by BL/KY/HV and the stock numbers. Two *Pcs* gene insertion lines, “*y*^*1*^, *w*^*67c23*^; *P{GSV2}pcs*^*GS7166*^*/SM1”* (KY201057) and “*w*; P{GawB}pcs*^*NP2623*^*/CyO”* (KY104263) were crossed to “*y, w, eyFLP; FRT42D”* flies and combined *FRT42D* with *pcs*^*NP2623*^ or *pcs*^*GS7166*^. The flies with *FRT42D, pcs*^*NP2623*^, or *pcs*^*GS7166*^ chromosomes were crossed to “*y, w, eyFLP; FRT42D, P3RFP”* to obtain mosaic eyes. To excise the *pcs*^*NP2623*^ or *pcs*^*GS7166*^ insertions, the fly lines with the *FRT42D* combined *pcs*^*NP2623*^ or *pcs*^*GS7166*^ chromosomes were crossed to “*w*; wg*^*Sp-1*^*/CyO; ry*^*506*^, *Sb*^*1*^, *P{ry[+t7.2]=Delta2-3}99B/TM6B, Tb*^*+*^*”* (KY107139). In the flies with white eyes, excisions were confirmed by sequencing. pUAS-V5::Pcs made here was crossed to Rh1-Gal4 and GMR-Gal4 to investigate the localization. *Rab11*^*EYFP*^ is a gift from Dr. Brankatschk (Max Planck Institute) (Dunst et al., 2015).

To target pcs gene, two CRISPR targets (ACACCCGGCTCGCGGGGTCCTGG and GTGCGACTATCCCTCCATAGCGG) on the exon4 of pcs gene were designed using CRISPR Optimal Target Finder (http://tools.flycrispr.molbio.wisc.edu/targetFinder/) (Gratz et al., 2014). BbsI digested fragment containing gRNA core and dU6:3 promoter was PCR amplified from pCFD4-U6:1_U6:3-tandemgRNAs (gift from Simon Bullock (Addgene plasmid # 49411) (Port et al., 2014), with primers pcs-g3-F1(5’-GTGCGACTATCCCTCCATAGGTTTTAGAGCTAGAAATAGCAAGTTAAAATAAGG-3’) and pcs-g2-R1(5’-GGACCCCGCGAGCCGGGTGTCGACgttaaattgaaaataggtctatatatacgaac-3’) and then pcs-g3-F2(5’-ggagggaggGAAGACccTTCGTGCGACTATCCCTCCATAGGT-3’) and pcs-g2-R2(5’-ccacccaccGAAGACccAAACGGACCCCGCGAGCCGGGTGTCG-3’), digested with BbsI, and cloned between BbsI sites of pCFD4-U6:1_U6:3-tandemgRNAs, resulting a phi31-transformation vector Pcs-CR1, which encodes the two guide RNAs targeting pcs gene. The plasmid pCFD4-Pcs was injected into embryos of PBac{yellow[+]-attP-3B}VK00033 (Bloomington stock center #9750) by BestGene Inc. (Chino Hills, CA, USA) to generate transgenic lines carring gRNA on the third chromosome.

*w; FRT42D; Pcs guide RNA / Vasa-Cas9 or Nos-Cas9* were crossed with “*y, w, eyFLP; Sp, heat-shock Hid / CyOGFP”* and the male progeny, *y, w, eyFLP; * 42D / CyOGFP* is crossed with “*y, w, eyFLP; Sp, heat-shock Hid / CyOGFP”* to make stocks. Males of each stock lines are crossed with *y, w, eyFLP; RFP42D; Arrestin2GFP* and observed Arrestin2GFP to judge the localization of Rh1. We observed 42 stocks and obtained three independent stocks, *Pcs*^*n1*^, *Pcs*^*n2*^ and *Pcs*^*Δ1*^ with four, two and 121 nucleotide deletions in Pcs coding region. In *Pcs*^*n1*^ allele, PcsS273stop476 is changed to 38amino acids (LRAARCLWAQKLPKQLPKLKMRKMPATTMKREPGSSEV*), in *Pcs*^*n2*^ allele, PcsI274stop476 is changed to 18amino acids (SGQPDVFGRKNSPSSCRN*) and in *Pcs*^*Δ1*^ allele, PcsP237stop476 is changed to 35amino acids (ARCLWAQKLPKQLPKLKMRKMPATTMKREPGSSEV*). Unfortunately, we lost *Pcs*^*n2*^ allele.

Insertions in two *TRAPP* genes, *PBac{PB}TRAPP2C*^*c00766*^*”* (HVc00766) and *Mi{ET1}TRAPPC11*^*MB06920*^ (BL25338) were combined with *FRT80B*. TRAPPC6 insertion, *TRAPPC6*^*f00985*^ (BL18399) was combined *FRT82B.* TRAPPC8 insertion line, “y^d2^ w^1118^ ey-FLP; P{lacW}l(3)76BDm(TRAPPC8)^L3809^, FRT80B/TM6B*”* was obtained from the Kyoto Drosophila Genomics and Genetic Resource (DGGR) (KY111022). To obtain mosaic eyes, the flies with *TRAPP2C*^*c00766*^, *FRT80B* or TRAPPC8^L3809*0*^, *FRT80B* chromosome were crossed to “*y, w, eyFLP; P3RFP, FRT80B”*, and the flies with *TRAPPC11*^*MB06920*^, *FRT80B* chromosome were crossed to “*y, w, eyFLP, FRT80B”* as *TRAPPC11*^*MB06920*^ hovers GFP. The flies with *FRT82B, TRAPPC6*^*f00985*^ were crossed to “*y, w, eyFLP; FRT82B, P3RFP”* to obtain mosaic eyes. We obtained the deletion alleles, *TRAPPC9*^*3D12*^ and *TRAPPC10*^*10F4*^ by imprecise excisions of the insertions in the 5’UTR, *TRAPPC9*^*KG04460*^ (BL13600) and in promoter, *TRAPPC10*^*EY00704*^ (BL15036). TRAPPC9^3D12^ or TRAPPC10^10F4^ has a deletion from 20654257-20656299 or 15217094-15219239.

### Fly retina immunostaining

Fixation and staining were performed as described previously (Satoh and Ready, 2005). Primary antisera were as follows: rabbit anti-Rh1 (1:1000) (Satoh et al., 2005), mouse monoclonal anti-Na^+^K^+^ATPase α subunit (1:300 ascite) (Developmental Studies Hybridoma Bank (DSHB), IA, USA), mouse monoclonal anti-Chp (24B10) (1:15 supernatant) (DSHB), rabbit anti-TRP (1:2000) (a gift from Dr. Montell, Johns Hopkins University), rat monoclonal anti DE-Cad (1:20 supernatant) (DSHB), rabbit anti-GM130 (1:300) (Abcam #ab30637, Cambridge, UK), rabbit anti-MPPE (1:1000) (a gift from Dr. Han, Southeast University, Nanjing, China), rat monoclonal anti-p120 (1:12) (a gift from Dr. Goto, Rikkyo University) (Yamamoto-Hino et al., 2012), guinea pig anti-Rab6 (1:300) (Iwanami et al., 2016), mouse anti-V5 monoclonal antibody: 6F5 (1:150) (WAKO Chemical #CTN3094, Osaka, Japan), rabbit anti-V5 (1:300) (MBL #PM003, Nagoya. Japan), and rat anti-Rab11 (1:250) (made here). Secondary antibodies were anti-mouse, anti-rabbit, anti-rat, and/or anti-guinea pig antibodies labelled with Alexa Fluor 488, 568, and 647 (1:300) (Life Technologies, Carlsbad, CA, USA). Images of samples were recorded using an FV1000 confocal microscope (60× 1.42 NA objective lens; Olympus, Tokyo, Japan). To minimize bleed-through, each signal in double- or triple-stained samples was imaged sequentially. Images were processed in accordance with the Guidelines for Proper Digital Image Handling using Fiji, Affinity photo, and/or Adobe Photoshop CS3 (Adobe, San Jose, CA, USA). The quantification of the intensity of Rh1 or Rab11staining in photoreceptor cytoplasm, we used more than 3 mosaic retinas with more than 20 wild type and more than 20 mutant photoreceptors in each retina. Area of cytoplasm or whole cells and also their staining intensities were measured by Fiji (Schindelin et al., 2012).

### S2 cell immunostaining

S2 cells expressing myc::APEX2::Pcs were fixed in 4% paraformaldehyde in 1x PBS for 1 hr on ice. Cells were rinsed three times for 2 minutes each in 1x PBS and then treated for 5 min in 1x PBS containing 0.1% Triton, followed by another three 2 minutes rinses in 1x PBS. Cells were incubated 2 hours in mouse monoclonal anti-myc 9E10C (1:15 supernatant) (DSHB) and guinea pig anti-Rab6 (1:150) (Iwanami et al., 2016) with 5% bovine serum in 1x PBS. After three rinses for 2 minutes each in 1x PBS, cells were incubated for 2 hours in anti-mouse antibodies labelled with Alexa Fluor 488 and anti-guinea pig antibodies labelled with Alexa Fluor 568 (1:150) (Life Technologies, Carlsbad, CA, USA). Imaging and data processing were the same for Fly retina immunostaining.

### Electron microscopy

Flies were reared in the dark and the retinal samples were fixed at the late pupal stage. To avoid light dependent Rh1 endocytosis, fixation was performed within 3 min of transferring the pupae to light. Electron microscopy was performed as described previously (Satoh et al., 1997). Samples were observed on a JEM1400 electron microscope (JEOL, Tokyo, Japan), and montage images were taken with a CCD camera system (JEOL). For the quantification of the area occupied by the vesicles in the cytoplasm, we used 5 photoreceptors of more than 3 retinas. The areas of vesicles and cytoplasm were measured by Fiji (Schindelin et al., 2012).

### Stably transformed S2 cells by myc::APEX2::Pcs

Synthesized DNA fragment K-miniSOG-Ap (GGGGATCTAGATCGGGGTACCGCCACCATGGAACAGAAGTTGATTTCCGAGGAAGGTCTGACTAGTGGAGGAGGAGGTTCTGGTGGTGGTATGGAGAAGAGCTTTGTTATCACAGATCCGAGGCTTCCAGACAACCCGATCATTTTTGCGTCCGATGGGTTCTTGGAACTGACAGAGTATTCAAGAGAAGAGATTTTGGGCCGAAACGGTAGGTTCTTGCAGGGTCCCGAAACCGACCAAGCGACGGTTCAGAAAATCAGGGACGCAATTAGGGATCAAAGAGAGATCACAGTGCAACTCATAAACTACACGAAAAGCGGCAAAAAATTCTGGAACCTTCTTCATCTTCAACCCATGAGGGACCAGAAGGGGGAACTCCAGTATTTTATCGGTGTTCAGTTGGACGGCGGTGGTGGAGGTTCTGGTGGCGGTGGCTCGAGTGAGCAAAAGCTCATTTCTGAAGAGGACTTGTAAGGGCCCTTCGAAGGTAAGCCT, IDT) was digested and cloned into KpnI-ApaI site of pMT-puro (gift from David Sabatini, Addgene plasmid #17923) to generate pMT-mSOG4m. A DNA fragment encoding APEX2 was amplified from pcDNA3-APEX2-NES (gift from Alice Ting, Addgene plasmid # 49386) using primers GL3N2-APEX2 (5’-ggaggttctggtggtggtGCGGCCGCcGGAAAGTCTTACCCAACTGTGA-3’) and APEX2-AGL4 (5’-ACCAGAACCTCCACCACCaGGCGCGCCGGCATCAGCAAACCCAAGCT-3’) then with primers Sp-GL3 (5’-CTGACTAGTggaggaggaggttctggtggtggt-3’) and GL4-Xh(5’-CTCACTCGAGCCaccgccACCAGAACCTCCACCACC-3’), digested and cloned into SpeI and XhoI site of pMT-mSOG4m to generate pMT-mAPEX2m. Gene fragment of pcs was amplified from cDNA of w1118 third instar larvae, using pcs-GF2 (5’-GGATGTGTCTGTGTAGCAACGAG-3’) and pcs-GR2 (5’-TGATATGGGGCTGGCTGAAGAAGTC-3’) as primers. To construct pMK-V5-pcs, the coding region of pcs amplified with pcs-MK-Sp (5’-gatcttcatggtcgactagaATTATTCCAGAGAGCGCCTACGCAG—3’) and GL3-pcs-F (5’-ggaggaggttctggtggtggtTCGAGTGCAGAAGACGGCGAG-3’) together with N-terminal V5-epitope, were cloned into Kpn-ApaI site of pMK33-CFH-BD using Gibson assembly.

The coding region was amplified from pMK-V5-pcs, using msXh-pcs-F (5’-gaggttctggtggcggtGGCTCGAGTGCAGAAGACGGCGAG-3’) and msAp-pcs-R (5’-AGGCTTACCttcgaaGGGCCTTATTCCAGAGAGCGCCTACGCAGC-3’) and cloned into XhoI-ApaI site of pMT-mAPEX2m to obtain pMT-mAPEX2-pcs. To produce stable transformants, 0.5ml of S2 cells were transfected with 1µg of DNA using 3µl of FuGENE HD (Progema), and selected with 2µg/ml Puromycin and/or 200mg/ml Hygromycin B for 2-3 weeks.

*Drosophila* S2 cells (gifted from Dr. Gota Goshima at Nagoya University, Japan) were stably transformed with pMK-myc::APEX::Pcs selected using puromycin.

### EM imaging of myc::APEX2::Pcs

Expression of myc::APEX2::Pcs was induced by adding 0.5 mM CuSO_4_. DAB staining was performed using the protocol described by Martell *et al*. (Martell et al., 2017). S2 cells expressing myc::APEX2::Pcs were fixed in 2% glutaraldehyde, 2% paraformaldehyde, and 2 mM CaCl_2_ in 0.1 mM cacodylate buffer (pH 7.4) for 1 hr on ice. Cells were rinsed five times for 2 min each in chilled buffer, then treated for 5 min in buffer containing 20 mM glycine to quench unreacted glutaraldehyde, and this was followed by another five 2-min rinses in chilled buffer. A freshly diluted solution of 0.5 mg/mL (1.4 mM) DAB (Sigma) in chilled buffer (DAB solution) was added to cells for 1 min, and then the solution was replaced by DAB solution with 10 mM H_2_O_2_ in chilled buffer and kept at room temperature for 3 min. Cells were rinsed 5 × 2 min with chilled buffer and then centrifuged at 100 g for 1-2 min. Cell pellets were mixed with pre-heated/melted 10% agarose in 0.1 M cacodylate buffer without CaCl_2_ (around 40-50C°) and cooled for solidification. Solidified agarose with cells were cut into 0.5- to 1-µm cubes. Post-fixation staining was performed with 2% (w/v) osmium tetroxide (Electron Microscopy Sciences) for 30 min in chilled buffer. Agarose cubes with cells were rinsed 5 × 2 min each in chilled distilled water and then placed in chilled 2% (w/v) uranyl acetate in ddH_2_O overnight. Agarose cubes with cells were dehydrated in a graded ethanol series (50%, 70%, 90%, 99.5%) for 5 min each time, and they were treated 2 times with dehydrated ethanol for 10 min. After treatment with propylene oxide 2 times for 10 min, agarose cubes with cells were infiltrated in EPON-812 (Electron Microscopy Sciences) using 1:1 (v/v) resin and propylene oxide for 3 hours and were then transferred into 100% resin and left to sit overnight. Agarose cubes with cells in EPON-812 were polymerized at 100 °C for 20 h. Embedded agarose cubes with cells were cut with a diamond knife into 70-nm sections and imaged on a JEM1400 transmission electron microscope (JEOL, Tokyo, Japan) operated at 80 kV, and montage images were taken with a CCD camera system (JEOL).

### Antisera against Rab1 and Rab11

We designed new rat anti-Rab1 and Rab11 antisera, as the mouse anti-Rab1 and Rab11 antisera were depleted (Satoh et al., 2005). The 6xHis-tagged Rab1 and Rab11 proteins were expressed in *E. coli* pG-KJE8/BL21 (TAKARA) at 23 °C using the previously created *pQE60-Rab11* vector (Li et al., 2007; Satoh et al., 2005) and purified in native conditions using Ni-NTA Agarose (QIAGEN, Hilden, Germany). To obtain antisera, 3 rats for each Rab proteins were immunized 6 times with 120 µg 6xHis-Rab1 or 80 µg 6xHis-Rab11 proteins. We designated the resultant antisera Rat1 to 3 anti-Rab1 and Rab11. New rat anti-Rab1 and anti-Rab11 antisera recognized endogenous Rab1 or Rab11 in immunoblots and exhibited staining patterns similar to those exhibited by the previous mouse anti-Rab1 and anti-Rab11 antisera in immunocytochemistry (Figures S1).

### Transgenic flies for UAS-V5::Pcs

V5 tag and Pcs genes were cloned into pUAST (Drosophila Genomics Resource Center, Bloomington, IN, USA) to construct pP{UAST-V5::Pcs} and injected into embryos by BestGene Inc. (Chino Hills, CA, USA) to generate transgenic lines.

### Immunoblotting

Immunoblotting was performed as described previously (Satoh et al., 1997). 3 rat anti-Rab1 and 3 rat anti-Rab11 (1:2000 concentrated supernatant) as primary antibodies and HRP-conjugated anti-rat IgG antibodies (1:20000, Life Technologies) as secondary antibodies. Signals were visualized using enhanced chemiluminescence (Clarity Western ECL Substrate; Bio-Rad, Hercules, CA, USA) and imaged using ChemiDoc XRS+ (Bio-Rad).

## Author Contributions

T. S. and A.S. designed the study; Y.O. performed most of the laboratory experiments and analyzed the data; N.N. performed EM imaging of myc::APEX2::Pcs; R. I made anti-Rab11 antibodies; T. S supervised the experiments for the molecular biology and S2 cells. A. S. supervised all of aspects of the project. T.S and A. S. wrote the paper with inputs and final approval from Y.O., N. N. and R. I.

## Ethics

Animal experimentation: All experiments were performed in accordance with the guidelines and approvals from the Hiroshima University Animal Care Committee (University of Toronto Protocol #20012022). The wild-type male mice, C57/ Bl6 strain (Charles River Laboratories), were used to produce antibodies. Animals were housed in the Faculty of Arts and Science Biosciences Facility (BSF) under a 12-h light: 12-h d cycle with 2-5 animals/cage.

## Acknowledgments

We thank Drs. U. Tepass, C. Montell, C. S. Goto, and Brankatschk for kindly providing fly stocks and reagents. We also thank the Bloomington Drosophila Stock Center and the Drosophila Genomics and Genetic Resource (Kyoto Institute of Technology) for fly stocks.

## Supplemental Information Captions

**Fig S1.**
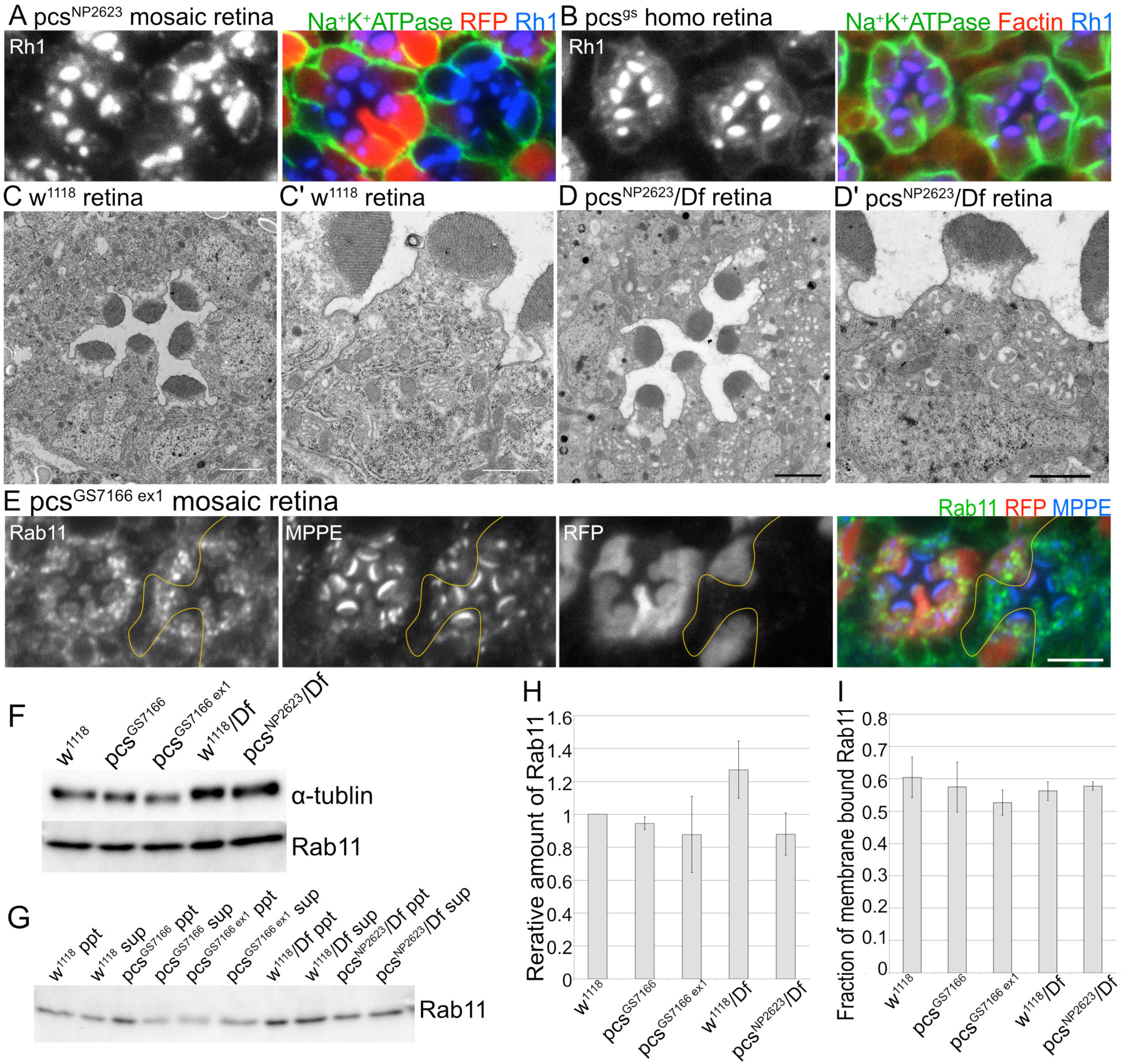
Rhabdomere proteins accumulate in the cytoplasm of *pcs*^*NP2623*^ homozygous photoreceptors. (A) Immunostaining of *pcs*^*NP2623*^ mosaic retina with anti-Na^+^K^+^ATPase (green) and anti-Rh1 (blue) antibodies. RFP (red) indicates wild-type cells. (B) Immunostaining of *pcs*^*gs*^ homozygous retina with anti-Na^+^K^+^ATPase (green) and anti-Rh1 (blue) antibodies. F-actin is visualized by phalloidin (red). (C, D) Electron micrographs of the ommatidia obtained from flies of the wild type, *w*^*1118*^ (C) and *pcs*^*NP2623*^/pcs deletion, Df2R2423 (D). Magnified electron micrographs are shown in C’ and D’. (E) Immunostaining of *pcs*^*GS7166 ex1*^ mosaic eye with anti-Rab11 (green) and anti-MPPE (Blue). RFP (red) indicates wild-type cells. (F) Immunoblotting of whole head extracts possessing the indicated genotypes with anti-α tubulin and anti-Rab11. (G) Immunoblotting of membrane and cytosolic fractions of head extracts with the indicated genotypes using anti-Rab11. (H) Relative amount of Rab11 in the whole-head extracts with the indicated genotypes. (I) Fraction of membrane-bound Rab11 in head extracts with the indicated genotypes. Scale bars: 5 µm (A, B, E, F), 2 µm (C, D), 1 µm (C’, D’).

**Fig S2.**
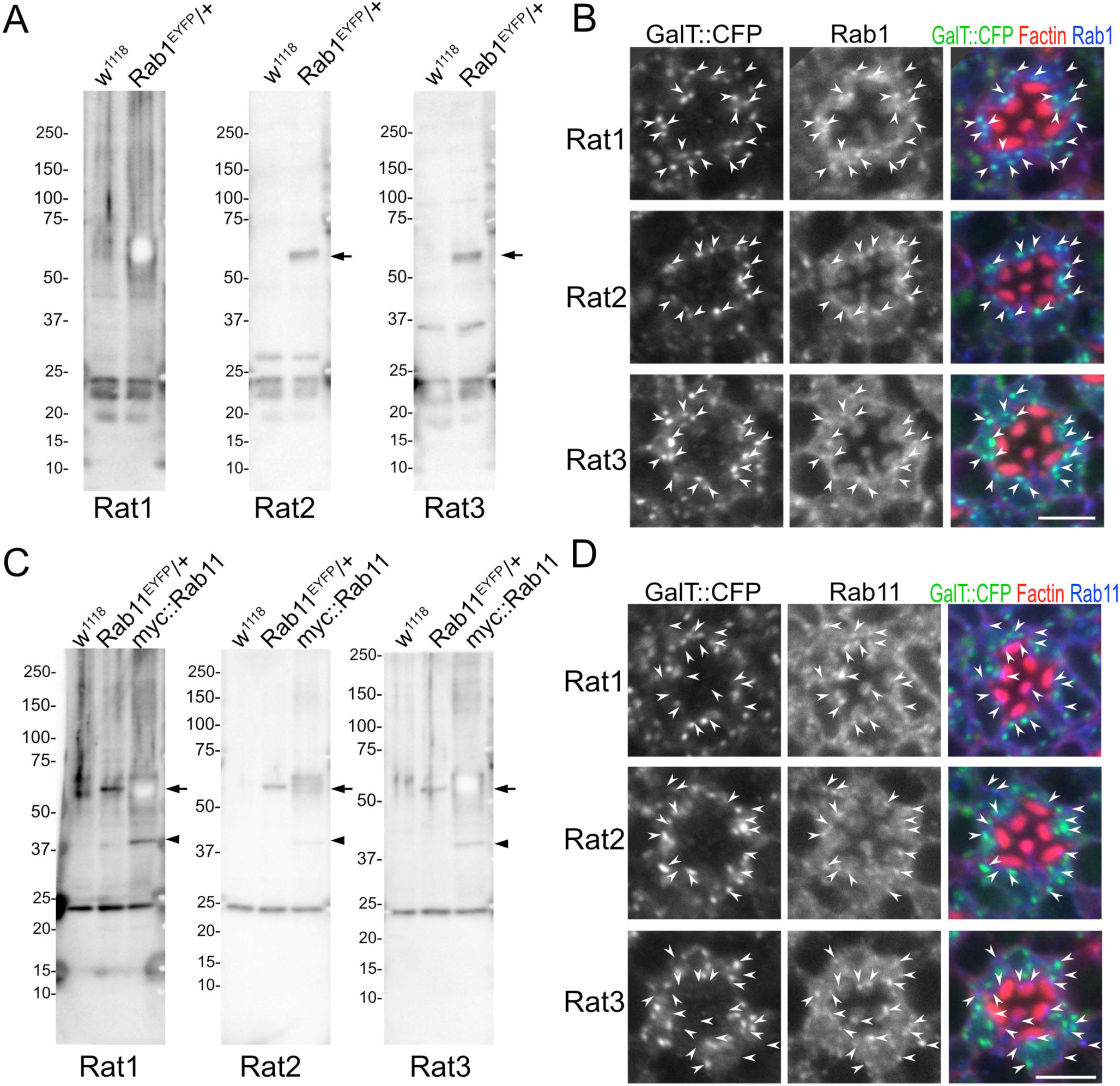
Verification of anti-Rab1 and anti-Rab11 antisera. (A, C) Immunoblotting of whole-head extracts with the indicated genotypes by rat anti-Rab1 (A) and anti-Rab11 (C) antiserum. All antisera against Rab1 and Rab11 recognized a band slightly below the band at approximately 25 kDa, which corresponds to the molecular weight of endogenous Rab1 and Rab11. All antisera against Rab1 recognized two additional bands that are smaller than the band near the 25 kDa marker, and these may be partially degraded products of Rab1. Arrows show EYFP::Rab1 or EYFP::Rab11, and arrowheads show 5x myc::Rab11. (B, D) Immunostaining of the retinas expressing GalT::CFP (green) using the anti-Rab1 (blue) (B) and anti-Rab11 (blue) (D) antisera. F-actin is visualized by phalloidin (red). Arrowheads show the co-localization of Rab1 or Rab11 and the Golgi marker, GalT::CFP. Scale bars: 5 µm (B).

**Fig S3.**
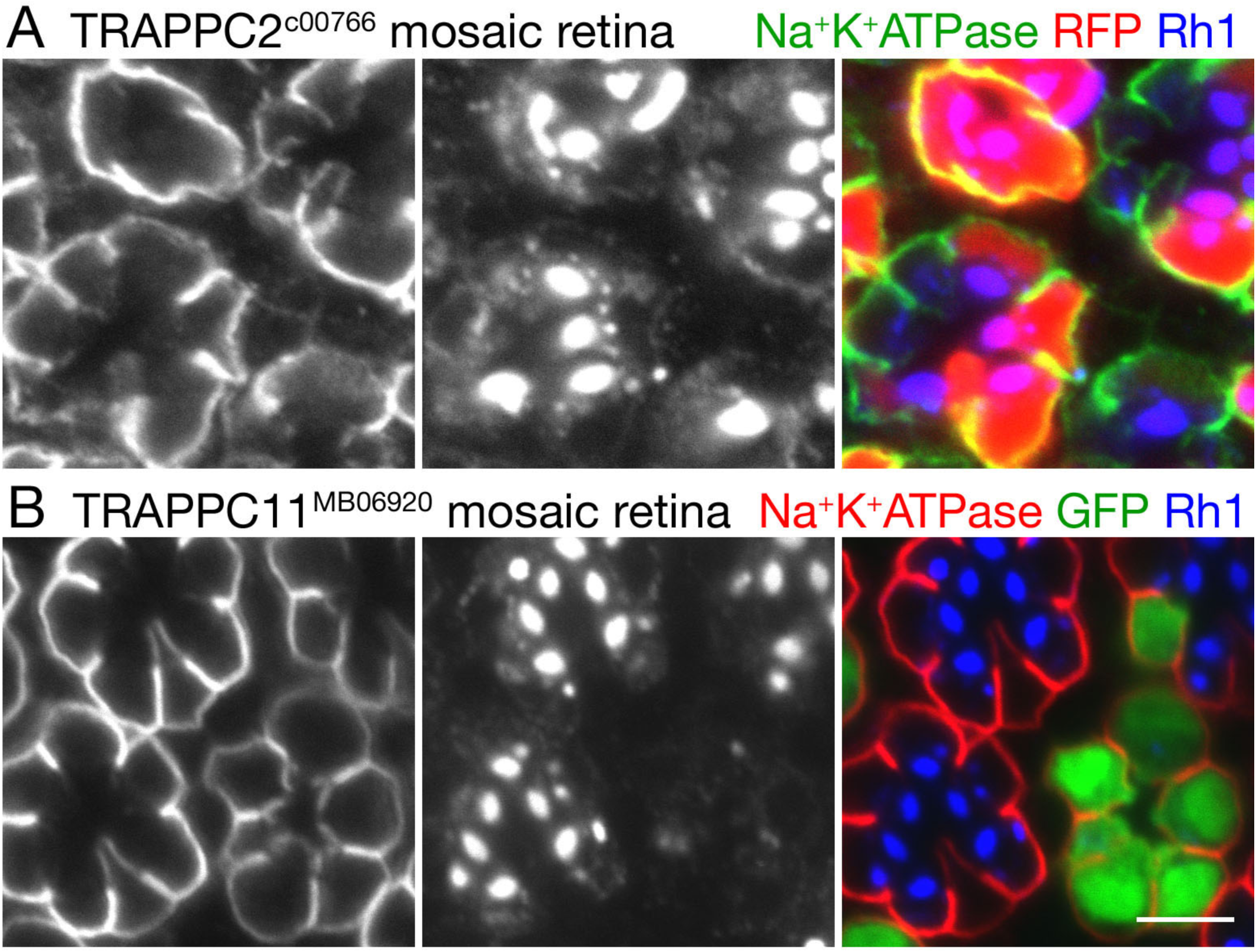
Rh1 transport is inhibited in *TRAPP2C*^*c00766*^ and *TRAPPC11*^*MB06920*^ homozygous photoreceptors. (A) Immunostaining of *TRAPP2C*^*c00766*^ mosaic eye with anti-Na^+^K^+^ATPase (green) and anti-Rh1 (blue) antibodies. RFP (red) indicates wild-type cells. (B) Immunostaining of *TRAPPC11*^*MB06920*^ mosaic eye with anti-Na^+^K^+^ATPase (red) and anti-Rh1 (blue) antibodies. GFP (green) indicates wild-type cells. Scale bars: 5 µm.

